# Larval densities of the protected Striped lychnis moth *Shargacucullia lychnis* (Lepidoptera: Noctuidae) in Buckinghamshire

**DOI:** 10.1101/2021.11.25.470005

**Authors:** Juliano Morimoto, Lucy Kerr

## Abstract

Natural history information is essential for ecologically-relevant inferences about (adaptive) responses in organismal biology. Yet, natural history data can be difficult to obtain, particularly for the developmental stages of holometabolous insects. This gap can compromise our ability to design controlled experiments that provide useful understanding of insect responses to changing environments and precludes our ability to understand how natural populations may respond to unpredictable climatic changes in their natural environment. In this study, we collated data from previous reports from the Butterfly Conservation Upper Thames Branch on the larval population density of *Shargacucullia lychnis* (Lepidoptera: Noctuidae) in Buckinghamshire. In the UK, *S. lychnis* is a protected species, for which natural history information can be invaluable for its effective conservation. We report here that the natural range of larval densities observed for *S. lychnis* across locations and years is 0.001 to 6.417 larvae per spike. More importantly, *S. lychnis* larval density has overall declined from 1996 to 2020, which could support previous reports of a contraction in population range for this species. Overall, this study provides invaluable information about larval population density for an important protected Lepidopteran species of the UK.

## Introduction

Natural history knowledge is paramount for understanding the biology of organisms (Markow, 2015) and for successful conservation efforts (Bury, 2006; Drew, 2011). Natural history institutions have adopted new ways to broaden access to natural history collections in order to facilitate discoveries in the field (Blagoderov, Kitching, Livermore, Simonsen, & Smith, 2012). Unfortunately, natural history as discipline has fallen out of fashion and has given way to hypothesis-driven research (Anderson, 2017; Mazzocchi, 2015; Schmidly, 2005). As a result, what once was the pillar upon which biological knowledge was built has now been cast aside, with the need to be re-integrated into ecological research (Anderson, 2017; Travis, 2020).

Natural history information is particularly absent for the insect larval stages in spite of larvae’s key ecological and economic roles in our societies (Morimoto, 2020). The developmental environment experienced by the larvae plays a key role in shaping adult individual fitness and, consequently, population survival (Than, Ponton, & Morimoto, 2020). An important factor determining the quality of larval developmental environment is larval density, for which virtually no natural history information is available for across holometabolous insect species (Than et al., 2020). This is because in holometabolous insects, larval density modulates life histories and fitness [reviewed by (Than et al., 2020)], including responses to temperature that can be crucial for adaptation to climate change (Henry, Renault, & Colinet, 2018; Lushchak et al., 2019). In fact, only recently have we had a direct glimpse of the natural population densities experienced by *Drosophila melanogaster* larvae, a species that is arguably the most well-studied insect in the world (Markow, 2015; Morimoto & Pietras, 2020). This study revealed that the majority of laboratory methodologies might have failed to manipulate larval density in values that were ecologically relevant to understand responses to both low and high densities (Morimoto & Pietras, 2020). It is, therefore, not surprising that for other insects, little or no information on the natural history of larval density is available in the published literature. This lack of natural history information about larval density (and larval ecology more generally) precludes our broader understanding of ecological forces acting upon the early stages of insect life.

Here, we contribute to the field of developmental ecology by reporting the natural history description of the larval density of the Striped lychnis moth *Shargacucullia lychnis* Rambur (1833) (Lepidoptera: Noctuidae) in a region in the UK (Fig 1). In particular, we collated data on the number of *S. lychnis* larvae in their host plant, *Verbascum nigrum*, from surveys and reports from 1996 and 2020 generated by the Upper Thames Branch of the UK Butterfly Conservation. This allowed us to use open monitoring data to gain natural history knowledge of *S. lychnis* larval densities. In the UK, *S. lychnis* is primarily restricted to two regions in central southern England: one around Oxfordshire, Berkshire, and Buckinghamshire, and the other in Hampshire and West Sussex, although in the past the population range was wider, reaching Gloucestershire in the west to Essex in the east (Heath & Emmet, 1983; Waring & Townsend, 2017). This suggests that *S. lychnis* population might have undergone range contraction. As a result of *S. lychnis* populations trends and scarcity in the UK entomofauna, *S. lychnis* has been listed in the UK Biodiversity Action Plan (UK BAP), which is a list of species deemed to be the most threatened and in need of conservation. Thus, the knowledge of larval densities can aid future conservation efforts, both by serving as a proxy for population size and dynamics (Fahrig & Paloheimo, 1988; Scott, 1994) but also by informing the species-specific patterns of competition that may modulate responses to anthropogenic pressures across the range of the species distribution.

**Figure 1.**
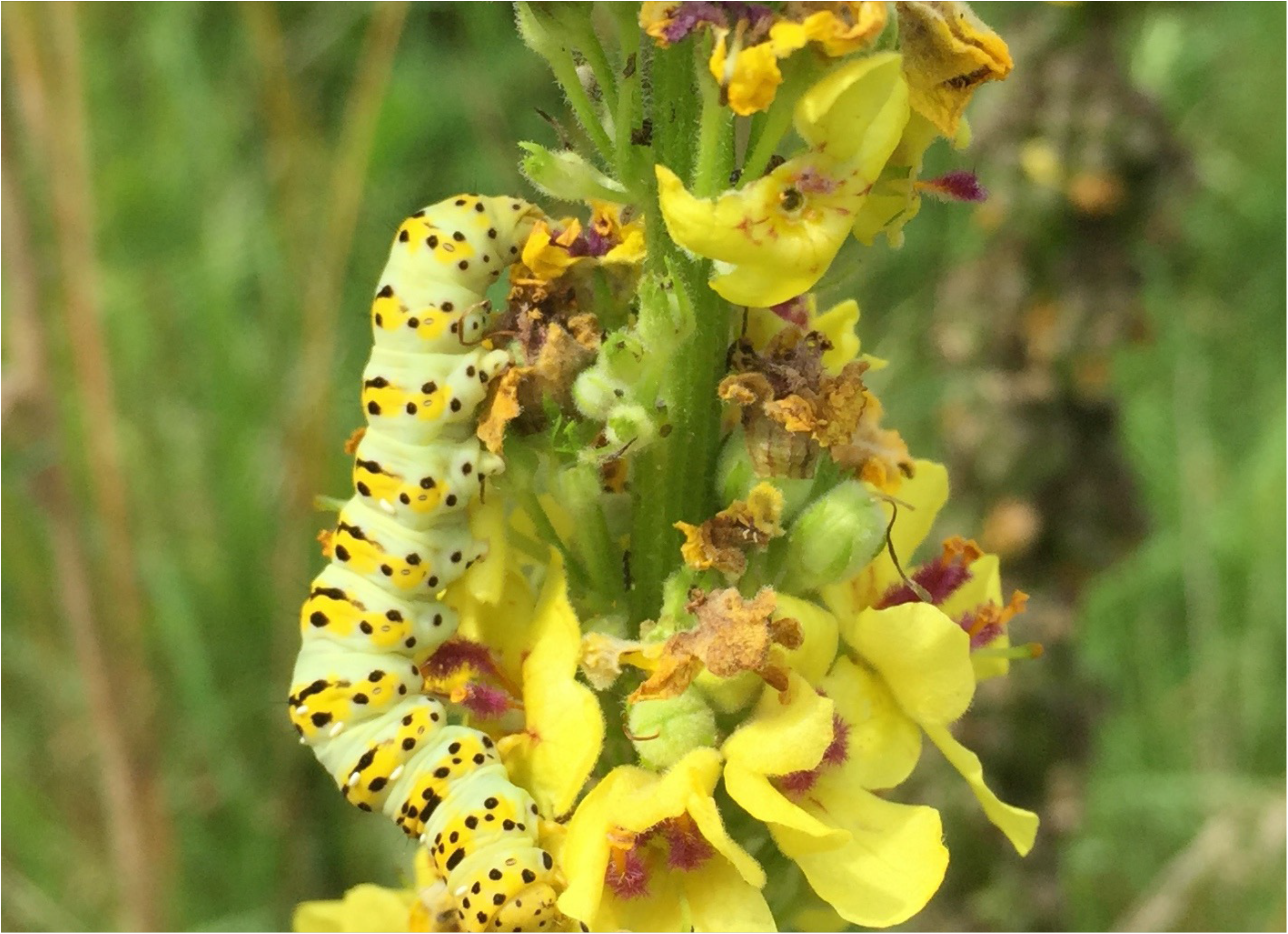
*S. lychnis* larvae feeding on a spike of the host plant (photo by Peter Cuss).

## Material and Methods

### Observations

All observations occurred in Buckinghamshire and were performed by the staff and volunteers of the Butterfly Conservation Upper Thames Branch. The observation methodology described here was provided by Peter Cuss (Peter Cuss, *pers. comm*.). Briefly, the number of spikes per site and the number of larvae was observed directly by the observers. Location determined the exact methods through which observations were made. For instance, at Homefield Wood where the host plant grows linearly along a bridleway, observations could be conducted by a single observer walking the route. Likewise, the Stoner to Henley site where there are few patches of hostplant along the route, a single observer could perform the observations. Larger sites, such as Green Farm, Bradenham and Watlington Hill, a team of volunteers assisted in the observations. Usually, volunteers line up along the field edge and each volunteer select a couple of trees or bushes in the distance as a reference point to form a corridor of ca. 20ft wide. Next, volunteers move across the area simultaneously while counting the larvae and spikes along the way. Observations were carried out at different times during the day. However, personal (uncontrolled) observations from the coordinator of the project Mr Peter Cuss suggests that time of day does not affect the number of larvae per spikes.

### Data collection and statistical analyses

All analyses were conducted in R version 3.6.2 (R Core Team, 2019). Data on the number of spikes of *Verbascum nigrum* and the number of *S. lychnis* caterpillars were collated from previously published reports generated by the Upper Thames Branch of the UK Butterfly Conservation. In total, we compiled a list of 55 survey locations from 1996 and 2020; at the time of the writing of the manuscript, data for 2019 and 2020 was not formally published as a report and was kindly provided by Peter Cuss. Sixteen locations were uniquely surveyed in 2019 and/or 2020, while the remaining locations were surveyed multiple times over the years. We crossed checked the data with earlier reports through visual inspection; no inconsistency in the reported data was found. As a result, we maintained the data entry from the most recent reports. We calculated an estimate of larval density by dividing the number of larvae by the number of spikes observed (i.e., larvae per spike) per location per year. In some cases, either the number of larvae or the number of spikes was missing (18%), which precluded a proper analysis of the larval density or the number of larvae was zero (11%). We therefore removed these data from the final analysis of larval density. In the end, 71% of the compiled data provided useful information about the distribution of *S. lychnis* larval density in nature (Fig 2). To assess whether larval density changed over time and across locations, we fitted a linear mixed model using the ‘lme4’ and ‘lmerTest’ packages (Bates, Sarkar, Bates, & Matrix, 2007; Kuznetsova, Brockhoff, & Christensen, 2017) with larvae per spike as dependent variable, observer’s name as random variable, and year and location as main effect independent variables. Data plots were made using the ‘ggplot2’ package (Wickham, 2016). Raw data is available as supplementary information.

**Figure 2.**
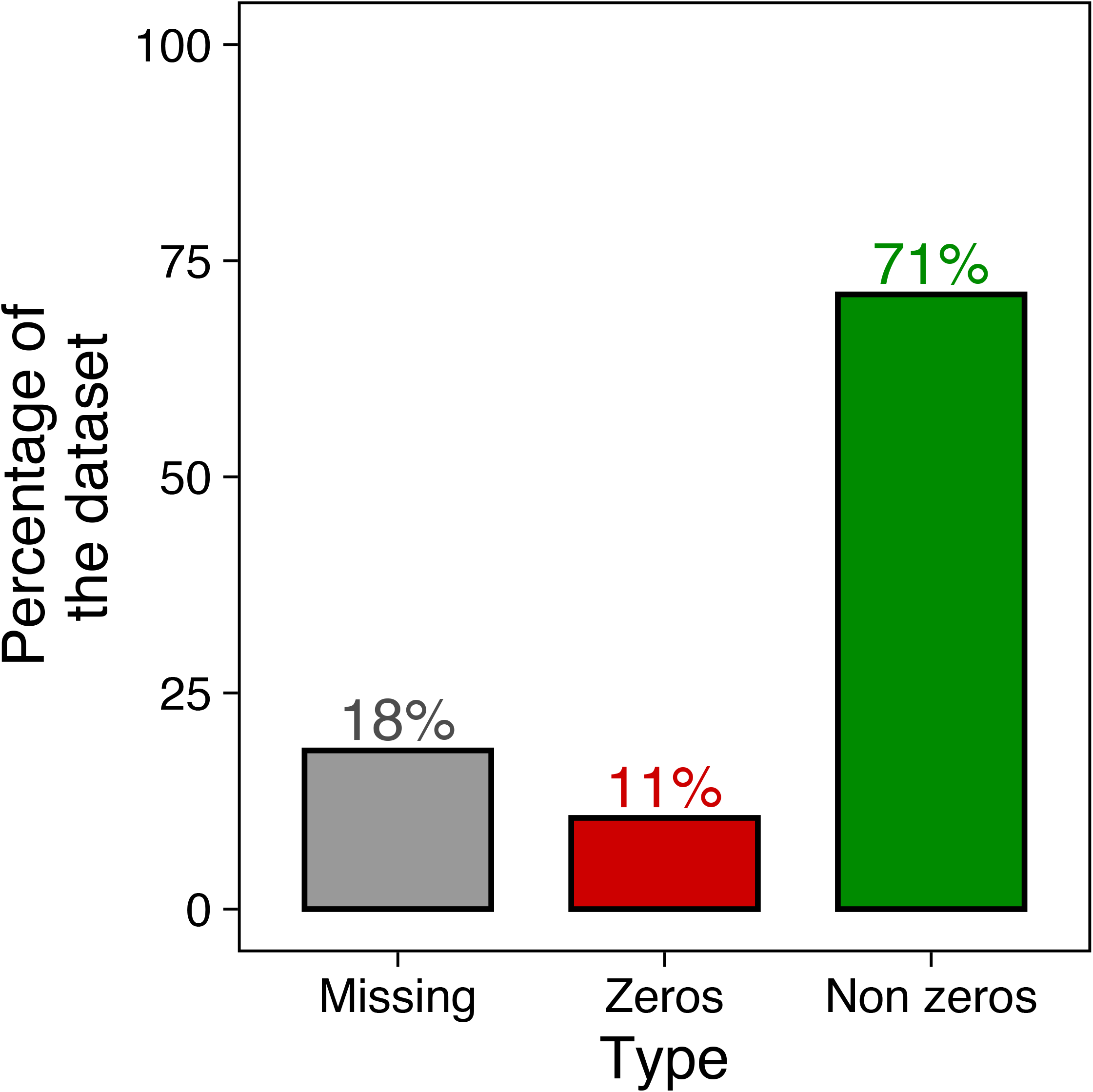
The proportion of missing, zero, and non-zero (usable) data compiled from the surveys.

## Results and discussion

*Shargacucullia lychnis* larval density distribution was right-skewed, with overall more common larval densities of two larvae per spike or less (Fig 3a). Log-transformation was effective in generating a normal-like distribution (Fig 3b), which facilitated the implementation of the linear mixed models.

**Figure 3.**
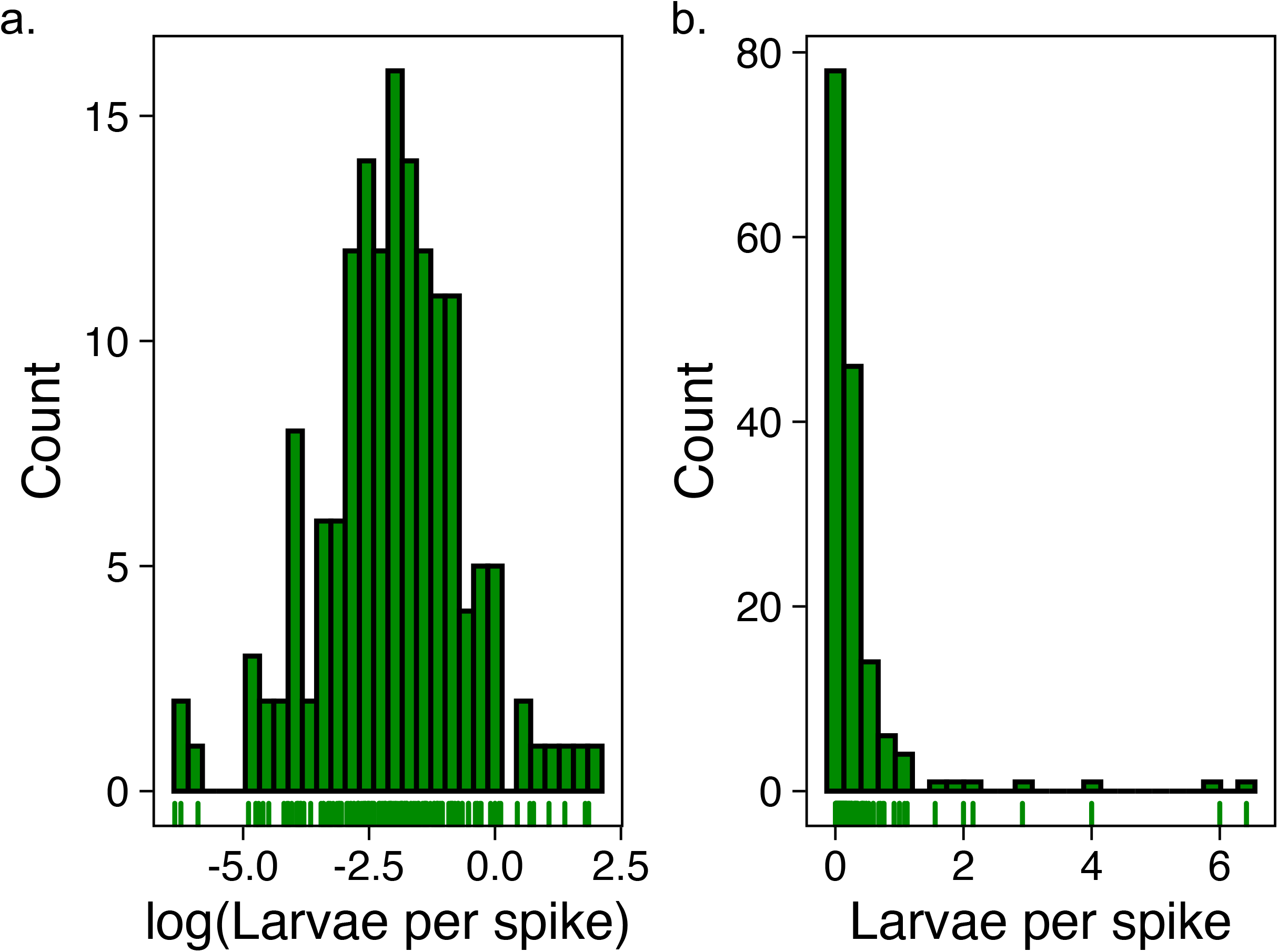
(a-b) The (log)distribution of larvae per spike observed in the datasets. Note that in (b), log-transformation resulted in a normal-distribution, which facilitated the implementation of the linear mixed models.

The regression model revealed that, overall, there has been a weak but significant decline in larvae per spike over the years (Est: -0.6174, std err: 0.1905; F_1, 100_ = 10.503; p = 0.001, Fig 4). Moreover, there was a statistically significant difference in larvae per spike across locations (F_53, 100_ = 1.952; p = 0.002) in which Bacombe Hill showed the lowest larvae per spike whereas Warburg displayed the highest larvae per spike. Importantly, the natural range of larval densities observed for *S. lychnis* across locations and years is 0.011 to 6.000 larvae per spike (Table 1). These results are significant because they help inform future the design of empirical studies conducted in controlled environments in order to disentangle physiological and behavioural responses to larval density in this and related species.

**Table 1.**
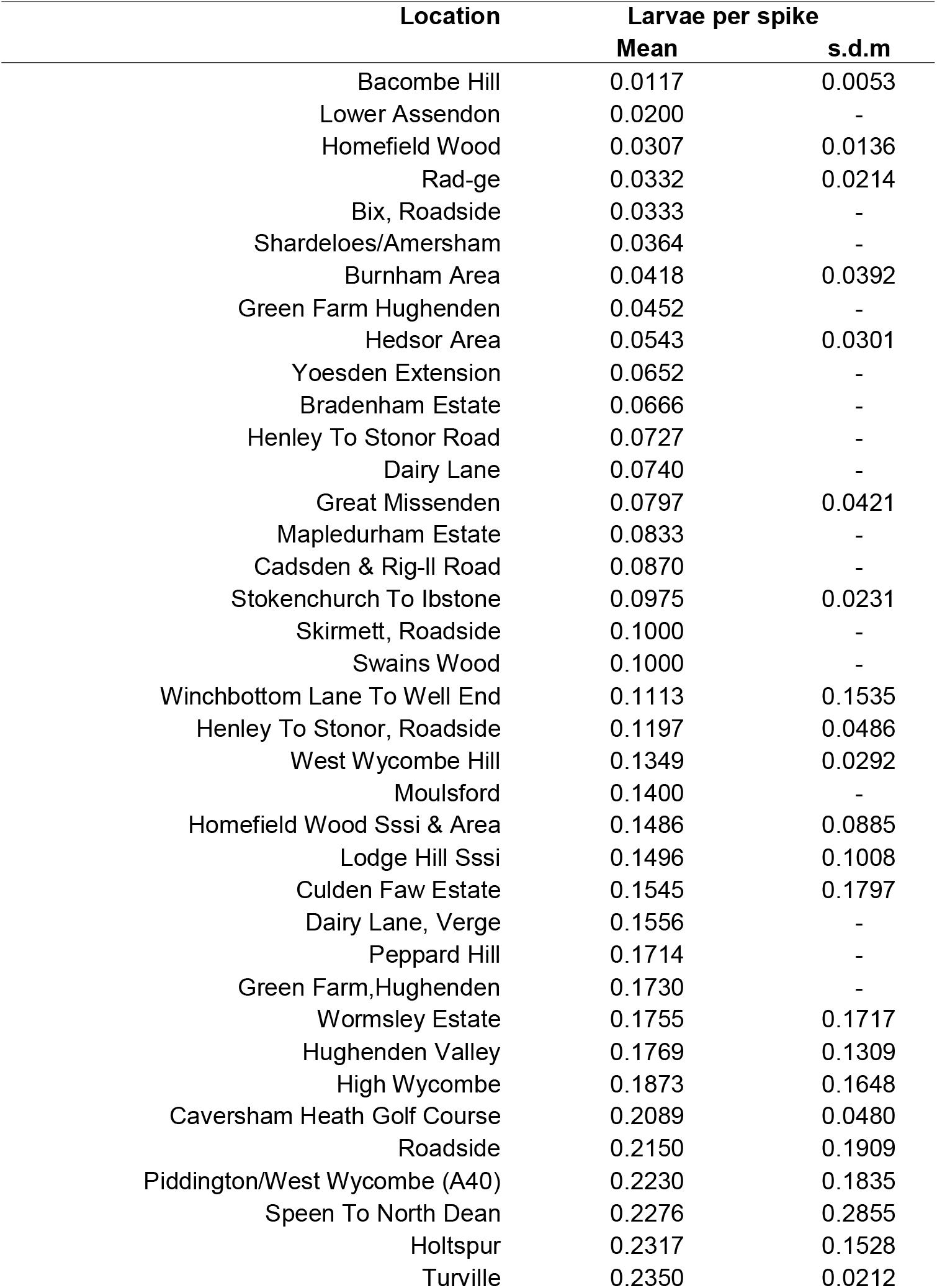
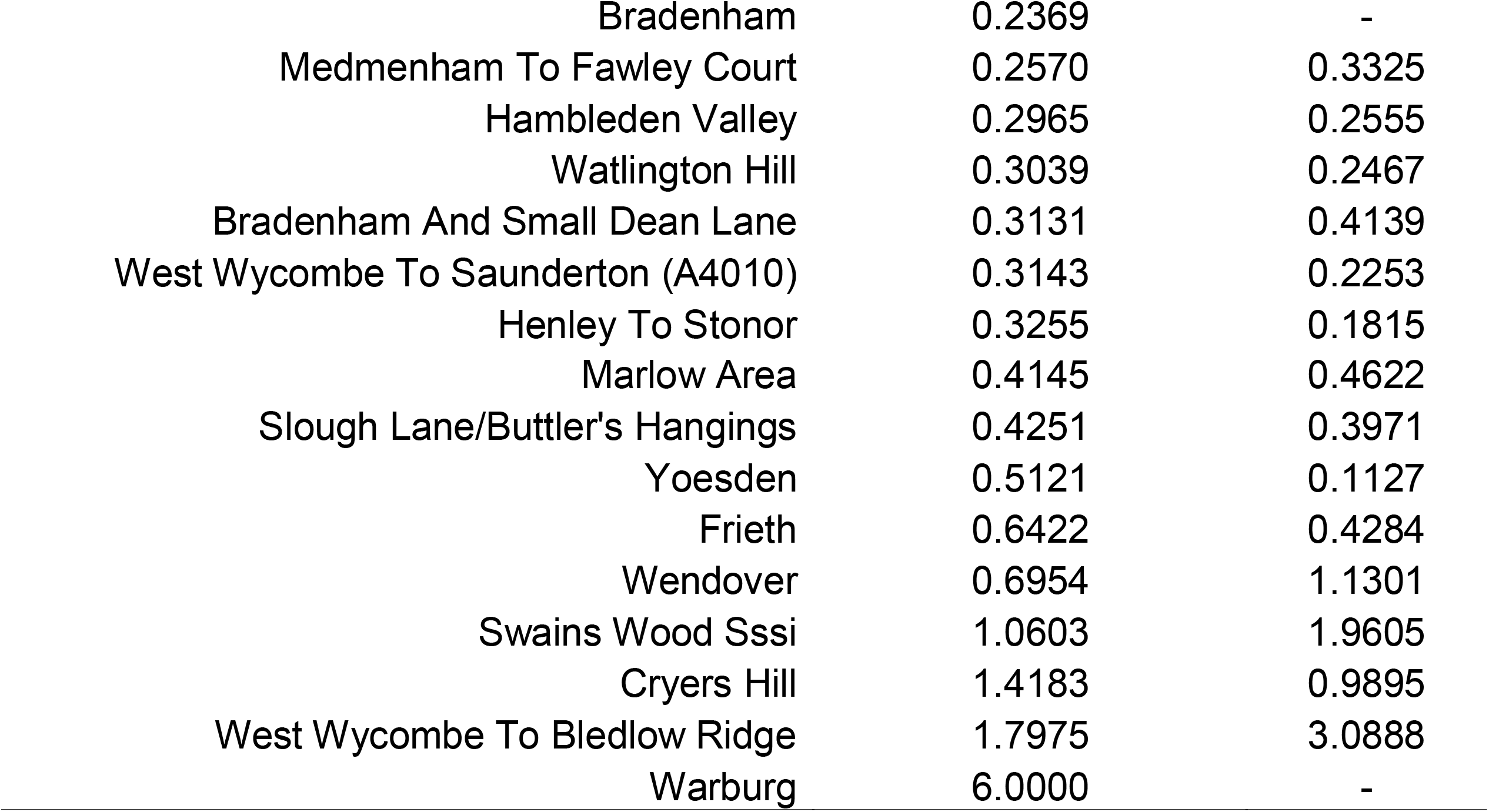
Summary of observed larvae per spike across locations. Cell entries that are absent for standard deviations of the mean (s.d.m) represent locations in which unique observations were made (see Methods).

**Figure 4.**
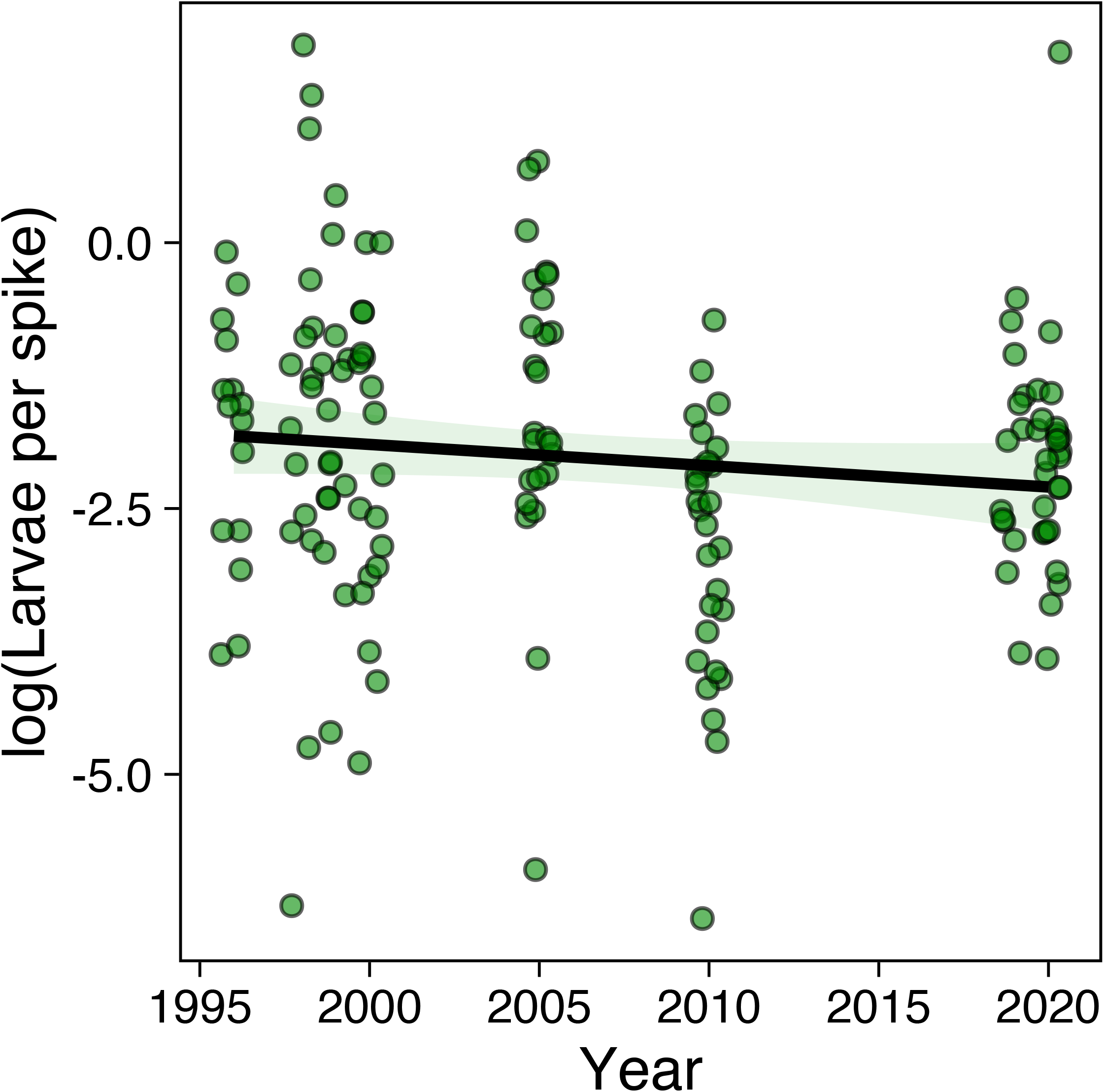
Larvae per spike over the years in which the monitoring occurred. Note that the y-axis is log-transformed.

As such, it aids ecologically meaningful insights into how the level of intraspecific competition at the larval stage of *S. lychnis* modulate life-history responses to other ecological factors that are relevant in our changing world, such as for example parasitism and temperature. Increased larval densities are associated with higher immunity in some Lepidopterans (‘density-dependent prophylaxis’) (Cotter, Hails, Cory, & Wilson, 2004). Climate change is expected to modulate between Lepidopterans and their parasitoids and thus understanding how larval density modulate immunity can help predict the implications to trophic interactions under different climatic conditions (Hufnagel & Kocsis, 2011). Likewise, larval density is known to interact with temperature to shape life-histories in other Lepidopterans, such as in *Chiasmia clathrata* (Lepidoptera: Geometridae) and the *Danaus plexippus* (Lepidoptera: Nymphalidae), helping our understanding of responses to warmer climatic conditions (Solensky & Larkin, 2003; Välimäki, Kivelä, & Mäenpää, 2013). Therefore, the natural history knowledge on the larval density of *S. lychnis* can be useful for future studies that investigate how *S. lychnis* may adapt to environments influenced by climate change. More broadly, the findings presented here provides a step towards addressing the knowledge gap in the field of developmental ecology, helping bridge the gap between current research in ecology and evolution with observational data from natural history (Travis, 2020).

## Acknowledgements

The authors would like to acknowledge Peter Cuss, Peter Hall, Sue Taylor, the Butterfly Conservation Upper Thames Branch, and all the volunteers who over the years have contributed to the collection of the data presented here. Without unprecedented efforts such as this, which provide invaluable information about the natural history of species, no studies by any academic would be relevant in the broader biological sense.

